# Phenotype-driven de novo molecular design from gene expression signatures

**DOI:** 10.64898/2026.07.21.739736

**Authors:** Yaxin Xu, Taojie Kuang, Shuang Ge, Haomin Wu, Mingqing Wang, Huan Xu, Fengwei An, Zhengyu Ma, Qiang Cheng, Zhixiang Ren

## Abstract

Target-based and structure-guided drug design remain central to modern drug discovery, but complementary strategies are needed when predefined targets or binding pockets do not fully capture disease biology. Gene-expression signatures provide scalable system-level readouts of disease and perturbation states, making them attractive inputs for phenotype-guided molecular design. However, preserving phenotypic information during molecular generation remains challenging, and chemically plausible molecules may lose connection to the intended biological response. Here, we present Tx2Mol, a transcriptome-guided framework that translates gene-expression signatures into candidate molecules while maintaining biological guidance throughout generation. We evaluated Tx2Mol across three biological settings: bulk gene perturbation, single-cell perturbation, and patient-derived disease signatures; and three validation dimensions: chemical plausibility, structural compatibility, and phenotypic preservation. Across 10 cancer-relevant bulk gene-perturbation benchmarks, Tx2Mol outperformed 9 transcriptome-guided baselines, improving maximum Tanimoto similarity to known ligands by 24.10% on average and by 50.67% on HDAC1. Structure-based analyses further supported structurally novel candidates with favorable predicted target binding. Tx2Mol also generalized to noisy single-cell perturbation profiles and preserved drug-induced transcriptional responses through in silico drug-perturbation validation. Patient-derived disease signatures further guided molecular generation toward approved-drug chemical space. Together, these results support gene-expression phenotypes as actionable guidance signals for phenotype-directed molecular design and candidate prioritization.

## 1 Introduction

Target-based and structure-guided drug design have made substantial progress and remain central strategies in modern drug discovery, particularly when disease-relevant targets and binding sites are well characterized. These approaches provide direct molecular interaction information and have been highly effective for rational ligand discovery. However, therapeutic response in complex diseases is also depend on pathway-level dysregulation, cellular state transitions, and context-dependent molecular networks [1, 2, 3]. When disease mechanisms remain incompletely understood, target or pocket information alone may not fully capture the biochemical response associated with disease modulation [4, 5]. These considerations motivate phenotype-guided molecular design as a complementary strategy that uses system-level biological readouts, rather than predefined target structures alone, as design inputs.

Gene-expression signatures represent one of the most informative and accessible phenotypic readouts for this purpose. As scalable and quantitative measurements of cellular state, they capture responses to disease processes, genetic perturbations, and drug treatment [6, 7]. By integrating signals across biological pathways and regulatory networks, transcriptomic profiles can bridge disease biology, perturbation responses, and molecular design [8, 9]. Nevertheless, effectively using gene-expression profiles for de novo molecular generation remains challenging. These profiles are high-dimensional, noisy, and strongly context dependent. The same perturbation can induce different transcriptional responses across cellular backgrounds, making it difficult to use gene expression as a direct molecular design signal [10, 11]. Thus, the central challenge lies not only in encoding complex transcriptomic profiles, but also in preserving their biological guidance throughout molecular generation [12, 13].

Prior studies have begun to connect transcriptomic phenotypes with molecular generation. Some approaches use gene-expression profiles for molecular retrieval, prioritization, or optimization of existing compounds [14, 15, 16, 17]. Others learn shared latent spaces between gene-expression signatures and molecules, enabling transcriptome-to-molecule sampling from learned joint representations [18, 19, 20, 21, 22]. More recent approaches adopt Transformer backbones or molecular language models to improve cross-modal representation learning [23, 24, 25, 26], as well as diffusion-based generation [27] and dual-channel designs [28, 29, 30]. Together, these studies demonstrate the feasibility of transcriptome-guided molecular generation. However, many existing methods use gene-expression profiles mainly as retrieval signals, optimization objectives, or global conditions for generation. Such one-shot or global conditioning may be insufficient to preserve biological guidance throughout molecular generation, where molecular cores, rings, functional groups, and peripheral structures are determined step by step. As a result, generated molecules may remain chemically plausible but only weakly linked to, or insufficiently validated against, the intended biological response. A key unmet need is therefore to maintain transcriptomic guidance throughout generation and to assess whether generated molecules preserve phenotype-relevant transcriptional responses.

Here, we introduce Tx2Mol, a transcriptome-guided post-training framework for de novo molecular generation. Tx2Mol aims to maintain effective gene-expression guidance during molecular generation through two complementary mechanisms. First, it converts gene-expression signatures and cell-line context into soft-prefix guidance tokens, making phenotypic information available throughout the molecular generation process. This allows the phenotype to influence both global molecular structure and local chemical features. Second, Tx2Mol employs bidirectional transcriptome–molecule contrastive alignment to strengthen the association between biological responses and their matched molecules. Together, these mechanisms enable the generation of chemically plausible molecules that are aligned with the input biological phenotype rather than defaulting to generic drug-like space.

We evaluated Tx2Mol across biological settings spanning controlled bulk gene-perturbation benchmarks, higher-resolution single-cell perturbation profiles, and patient-derived disease signatures. Across evaluation dimensions, we assessed whether generated molecules exhibit chemical plausibility, structural compatibility, and phenotypic preservation. In bulk gene-perturbation benchmarks, Tx2Mol improved hit-like recovery across cancer-relevant targets while maintaining molecular quality and novelty, with structure-based analyses further supporting plausible target engagement through docking and ligand efficiency. In single-cell settings, Tx2Mol generalized to noisy transcriptional profiles and preserved drug-induced transcriptional responses in closed-loop in silico analyses. Patient-derived disease signatures guided generation toward approved-drug chemical space while supporting broader therapeutic chemical exploration. Together, these results support gene-expression phenotypes as actionable guidance signals for phenotype-directed molecular design and candidate prioritization.

## 2 Results

### 2.1 Overview of Tx2Mol

To connect transcriptomic phenotypes with de novo molecular generation, we developed Tx2Mol, a transcriptome-guided post-training framework that maps gene expression signatures to molecular structures (Fig. 1). The key challenge is to maintain phenotype-relevant guidance throughout molecular generation. Gene-expression phenotypes are high-dimensional and context dependent, whereas molecular generation proceeds through local structural decisions over atoms, rings, and functional groups. If the phenotype is used only as an initial condition, this guidance can fade during generation, leading to chemically plausible but phenotype-agnostic molecules.

**Figure 1:**
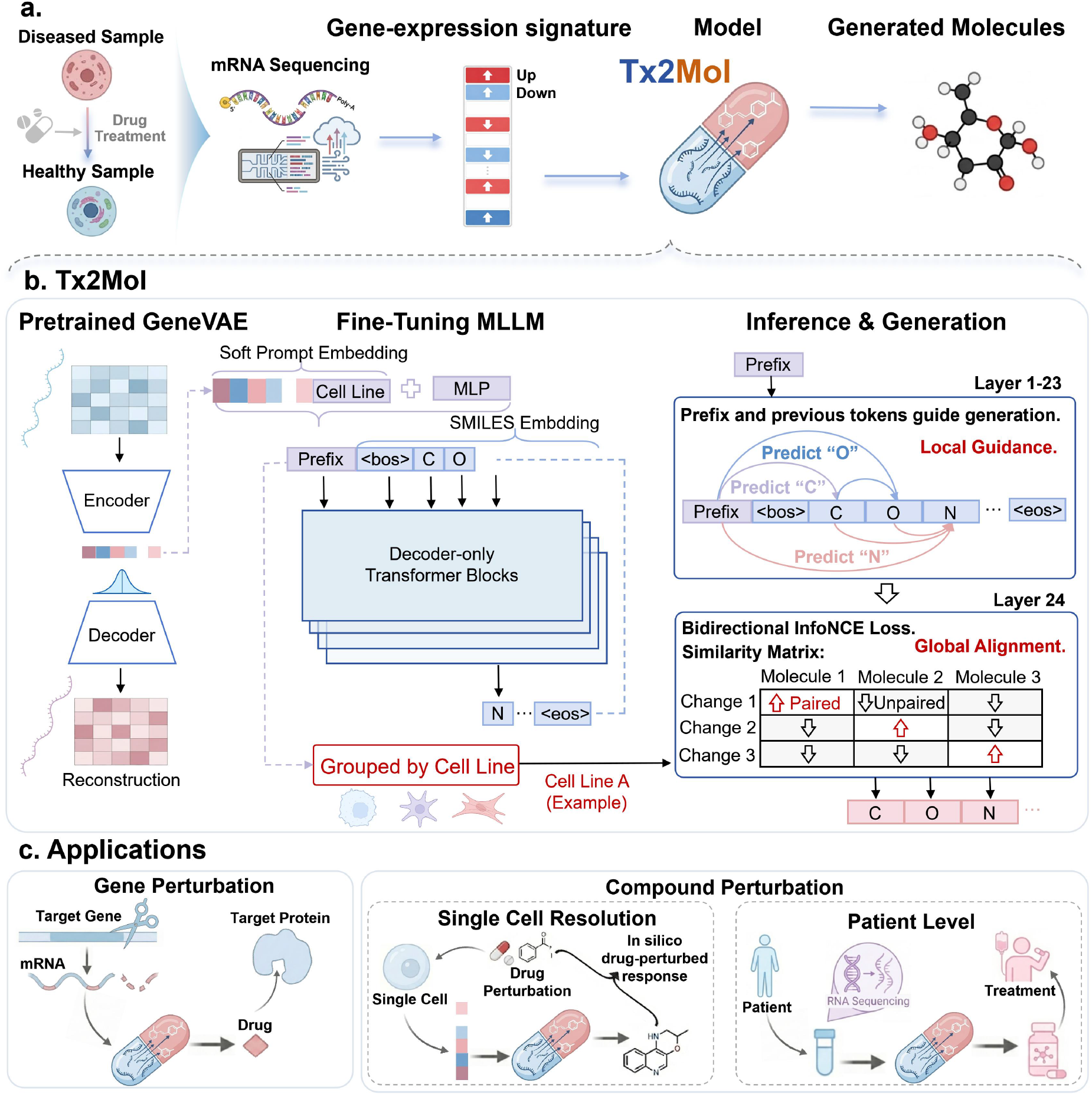
Overview of Tx2Mol for phenotype-guided molecular generation from gene expression signatures. **a**, Problem setup. Gene-expression changes derived from perturbation or disease states are used as phenotypic inputs to guide de novo molecular generation across bulk, single-cell, and patient-derived settings. **b**, Tx2Mol links gene-expression phenotypes with molecular generation through two complementary mechanisms. **Local guidance** converts gene-expression signatures and cell-line context into soft-prefix tokens that provide persistent phenotypic constraints during molecular generation. **Global alignment** uses bidirectional contrastive learning to align phenotypic responses with their matched molecules. **c**, Evaluation strategy across bulk gene perturbation, single-cell perturbation, and patient-derived disease settings, assessing hit-like generation, target relevance, generalization, transcriptional-response preservation, and therapeutic chemical-space guidance.

Tx2Mol was designed to address this problem. It first converts the gene-expression signature and cell-line context into soft-prefix tokens. These tokens allow each structural decision to access the phenotypic signal during generation. This makes the phenotype a persistent design constraint rather than a one-time input. Tx2Mol also uses bidirectional contrastive alignment to link each phenotypic response with its matched molecule. This encourages phenotype-relevant chemistry and reduces the tendency to generate generic drug-like molecules. Together, these mechanisms enable Tx2Mol to generate chemically plausible molecules that remain tied to the input biological state.

We evaluated Tx2Mol across bulk gene perturbation, single-cell perturbation, and patient-derived disease settings using a chemical-structural-phenotypic evaluation strategy (Fig. 1c). This strategy asks whether transcriptomic phenotypes can guide molecular generation beyond chemical plausibility alone. At the chemical level, we tested whether generated molecules recover hit-like and approved-drug-like chemical space while retaining molecular quality and structural novelty. At the structural level, we examined whether generated candidates show plausible target engagement using docking and ligand-efficiency analyses. At the phenotypic level, we evaluated whether generated molecules preserve the intended biological response through target specificity, single-cell compatibility, and in silico validation of drug-induced transcriptional responses. Together, these analyses connect transcriptomic guidance to molecular plausibility, target relevance, and transcriptional-response preservation.

### 2.2 Gene-perturbation phenotypes guide hit-like molecular generation

#### 2.2.1 Hit-like molecular generation with phenotype preservation

To evaluate whether gene-perturbation phenotypes can guide hit-like molecular generation, we benchmarked Tx2Mol on bulk perturbation signatures derived from 10 cancer-relevant targets (AKT1, AKT2, AURKB, CTSK, EGFR, HDAC1, MTOR, PIK3CA, SMAD3, and TP53). As shown in Fig. 2a, Tx2Mol achieves the highest maximum Tanimoto similarity to matched known ligands across all 10 targets, outperforming 9 transcriptome-guided generative baselines. Relative to the strongest baseline, Tx2Mol improves maximum Tanimoto similarity by 24.10% on average, including a 50.67% improvement on HDAC1. Detailed results for all targets are provided in Supplementary Table S1.

**Figure 2:**
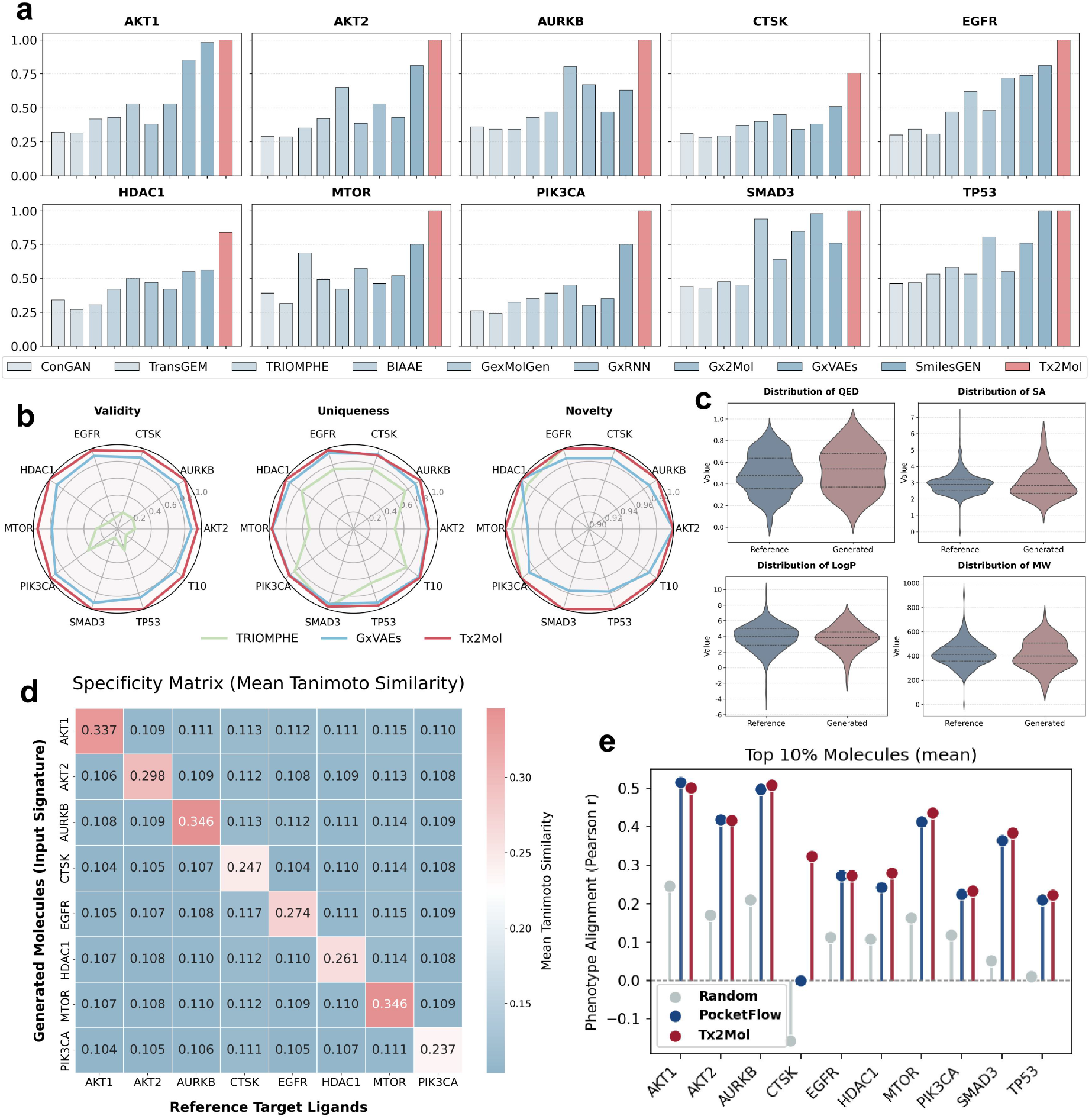
Bulk gene-perturbation phenotypes support hit-like molecular generation while preserving transcriptomic phenotypes. **a**, Comparison of Tx2Mol with 9 transcriptome-guided generative baselines on 10 cancer-relevant targets, evaluated by maximum Tanimoto similarity to matched known ligands. Tx2Mol performs best across all targets, with a 50.67% improvement on HDAC1 over the strongest baseline. **b**, Validity, uniqueness, and novelty relative to representative baselines, showing that improved hit-like recovery is achieved without compromising generation quality. **c**, Physicochemical property distributions of AKT1-generated molecules versus known ligands, showing improved QED and SA while maintaining comparable LogP and MW distributions. **d**, Specificity matrix of mean Tanimoto similarity between molecules generated from each perturbation signature and known ligands across targets, showing a clear diagonal enrichment and supporting transcriptomic phenotype specificity. **e**, Comparison of phenotype alignment for the top 10% molecules generated by Tx2Mol and PocketFlow across ten cancer-relevant targets. Random molecules are shown as the median reference.

Importantly, these gains do not compromise generation quality. Compared with representative baselines (TRIOMPHE and GxVAEs), Tx2Mol maintains competitive validity, uniqueness, and novelty (Fig. 2b). In addition, we further assessed the physicochemical properties of the generated molecules using Quantitative Estimate of Drug-likeness (QED), Synthetic Accessibility (SA), LogP, and Molecular Weight (MW). For AKT1, generated molecules show improved QED and SA while maintaining comparable LogP and MW distributions (Fig. 2c). Similar trends were observed across the remaining nine targets (Supplementary Fig. S4).

We next asked whether Tx2Mol preserved phenotype specificity rather than generating generic ligand-like molecules. To this end, we calculated the mean Tanimoto similarity between molecules generated for each perturbation signature and known ligands across all 10 targets. As shown in Fig. 2d, the resulting specificity matrix exhibits a clear diagonal enrichment, indicating that molecules generated from each perturbation signature are preferentially similar to the matched known-ligand set. Consistent results were obtained using cosine similarity, with significantly higher on-target than off-target similarity for all targets (*p <* 0.001; Supplementary Fig. S5). Together, these results show that Tx2Mol generates hit-like molecules while preserving transcriptomic phenotype specificity.

Chemical-level evaluations alone cannot establish whether generated molecules preserve the intended transcriptomic phenotype. We therefore trained an independent gene-expression predictor to estimate the transcriptomic responses induced by generated molecules. For each target, Tx2Mol and PocketFlow [31] generated 100 molecules from transcriptomic phenotypes and the corresponding protein pockets, respectively. We selected the top 10% molecules based on phenotype-alignment scores. Randomly sampled training-set molecules served as a reference. As shown in Fig. 2e, Tx2Mol consistently achieved stronger phenotype alignment across most targets. PocketFlow performed competitively on well-characterized targets such as AKT1, whereas Tx2Mol achieved substantially stronger phenotype alignment for CTSK, a more challenging target for pocket-based generation. These results provide orthogonal evidence that transcriptomic phenotypes directly guide the generation of biologically relevant molecules.

#### 2.2.2 Structure-based validation of generated candidates

Although the generated molecules exhibit favorable chemical characteristics and preserve the intended transcriptomic phenotype, it remains unclear whether these phenotypic responses arise from plausible interactions with the intended target proteins. We therefore performed structure-based validation using molecular docking with AutoDock Vina [32]. Five experimentally resolved target structures with well-defined binding pockets were selected for evaluation; detailed information is provided in Supplementary Table S2 and Fig. S6.

For each target, Tx2Mol generated 1,000 molecules for docking against the corresponding protein pocket. The top 10% ranked by Vina score were compared with the corresponding known ligands. As shown in Fig. 3a, the generated candidates consistently have lower Vina scores than the corresponding known ligands, indicating stronger predicted binding affinity. Because Vina scores systematically favor higher-MW molecules, we additionally report ligand efficiency (LE) to normalize predicted binding strength by molecular weight (Fig. 3b). Generated molecules show higher mean LE than known ligands across targets, indicating that improved docking is not simply driven by increased molecular weight.

**Figure 3:**
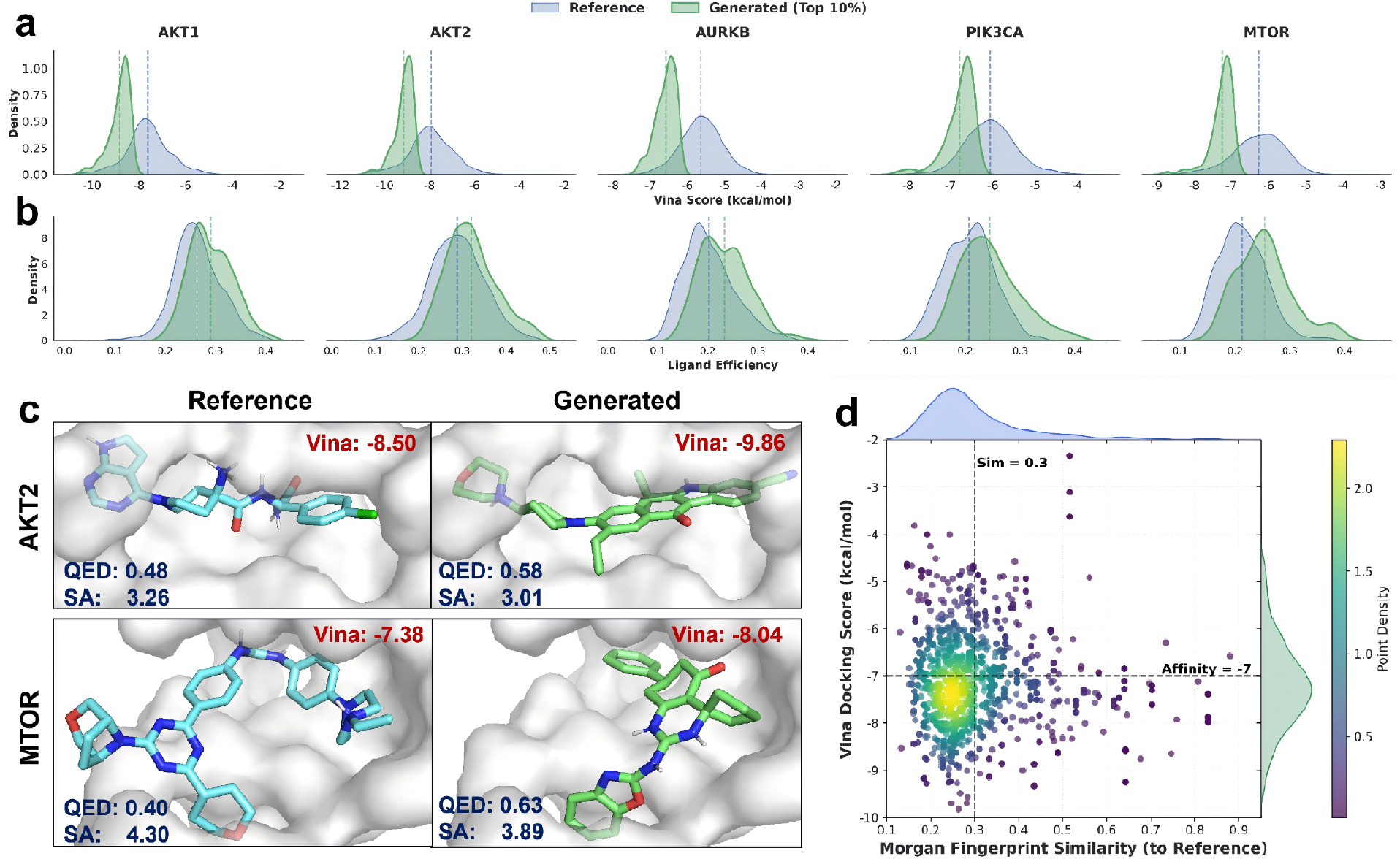
Structure-based analyses support target-relevant generation beyond 2D similarity. **a**, Vina score distributions for known ligands and the top 10% generated molecules across 5 targets, showing consistently stronger predicted binding affinity for the generated molecules. **b**, Ligand-efficiency distributions for the same molecules, controlling for the MW bias of Vina scores and showing that improved docking performance is not explained simply by larger molecules. **c**, Representative docking poses for AKT2 and MTOR, showing plausible binding modes together with favorable QED and SA. **d**, Joint analysis of 2D chemical similarity and 3D binding affinity for AKT1, showing that 37.57% of generated molecules achieve Vina scores below −7 kcal/mol despite Tanimoto similarity below 0.3.

In addition, representative docking poses for targets AKT2 and MTOR illustrate that Tx2Mol can recover plausible binding modes while maintaining favorable QED and SA (Fig. 3c). Finally, we jointly analyzed 2D chemical similarity and 3D binding affinity by comparing Morgan fingerprint similarity to known ligands with Vina docking scores. As shown for AKT1 in Fig. 3d, 37.57% of generated molecules have maximum Tanimoto similarity below 0.3 while achieving Vina scores below −7 kcal/mol. Consistent results were observed across the remaining four targets (Supplementary Fig. S7). These results indicate that Tx2Mol generates structurally distinct candidates beyond close analogs of known ligands. Together, these analyses extend the validation of Tx2Mol from chemical and transcriptomic phenotypes to target-level structural compatibility.

### 2.3 Single-cell perturbation phenotypes extend transcriptome-guided molecular generation

#### 2.3.1 Generalization to single-cell perturbation phenotypes

We next investigated whether Tx2Mol generalizes from bulk transcriptomic phenotypes to single-cell perturbation profiles, which provide higher-resolution yet substantially noisier transcriptional measurements. Using the sci-Plex3 [33] dataset, we extracted single-drug perturbation profiles (10 µM, 24 h) from A549, K562, and MCF7 cell lines. For each perturbation profile, Tx2Mol generated 100 candidate molecules, which were evaluated by maximum Tanimoto similarity to the corresponding perturbing compound.

Tx2Mol consistently outperforms ablated variants lacking soft-prefix guidance or InfoNCE alignment (Fig. 4a), demonstrating that transcriptomic phenotypes remain effective guidance signals for molecular generation at single-cell resolution. Removing soft-prefix guidance caused the largest performance drop, highlighting the importance of token-level conditioning during molecular generation. InfoNCE alignment further improved robustness across heterogeneous single-cell perturbation profiles.

**Figure 4:**
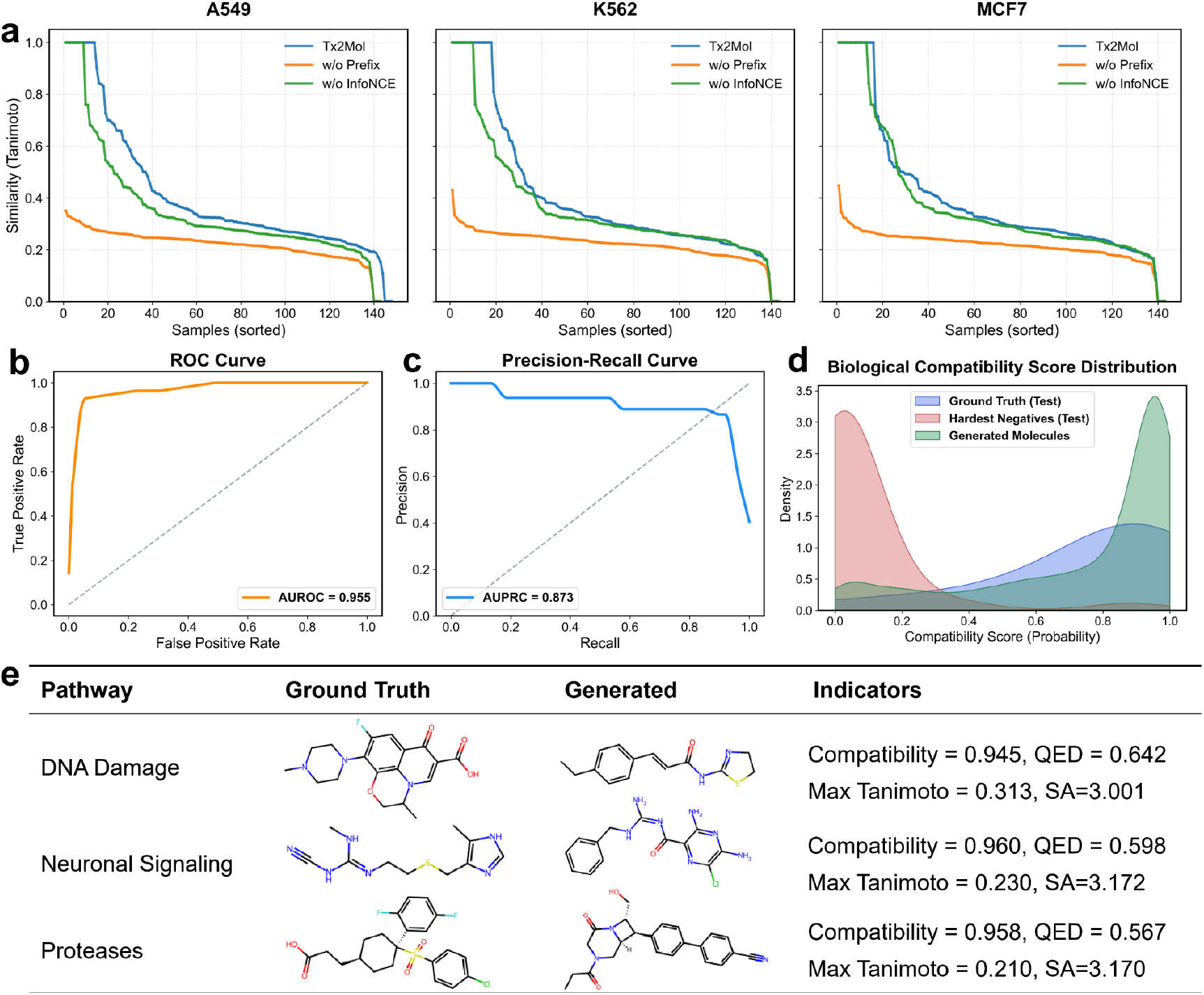
Single-cell perturbation phenotypes support the generalization of transcriptome-guided molecular generation. **a**, Evaluation on single-cell perturbation profiles from A549, K562, and MCF7 cell lines using maximum Tanimoto similarity to the matched perturbing compounds. Tx2Mol consistently outperforms ablated variants lacking soft-prefix guidance or InfoNCE alignment. **b–d**, Performance of a learned compound-phenotype compatibility scorer trained on true transcriptome–molecule pairs and hard negatives. **b**, ROC analysis. **c**, Precision-recall analysis. **d**, Compatibility-score distributions for true pairs, hard negatives, and Tx2Mol-generated molecules. **e**, Representative high-scoring molecules from the DNA Damage, Neuronal Signaling, and Proteases pathways, showing structurally distinct candidates with reasonable QED and SA.

We then asked whether the generated molecules preserve phenotype-level compatibility beyond chemical similarity. Directly testing hundreds of generated molecules experimentally is impractical. Inspired by GEMGen [25], we trained a compound–phenotype compatibility scorer on A549 cells. We treated true compound-signature pairs as positives and used the three most structurally dissimilar compounds as hard negatives. The scorer distinguished true pairs from hard negatives, achieving an AUROC of 0.955 (Fig. 4b) and an AUPRC of 0.873 (Fig. 4c), with high compatibility scores preferentially assigned to true pairs (Fig. 4d).

Representative examples from three pathways: DNA Damage, Neuronal Signaling, and Proteases, are shown in Fig. 4e. High-scoring molecules are often structurally distinct from the matched perturbing compounds while retaining reasonable QED and SA. For a given pathway perturbation signature, Tx2Mol therefore does not collapse to nearest-neighbor analogues, but instead generates diverse candidates spanning multiple chemical archetypes for the same pathway perturbation. Together, these results demonstrate the robust generalization of Tx2Mol from bulk to single-cell perturbation phenotypes while preserving phenotype compatibility and chemical diversity.

#### 2.3.2 In silico validation of preserved single-cell transcriptional responses

Having shown that Tx2Mol generalizes to single-cell perturbation phenotypes, we next asked whether the generated molecules preserve the transcriptional responses induced by the original drugs. To address this question, we used chemCPA [34] as an independent in silico perturbation predictor. This enabled closed-loop phenotypic validation from input expression to Tx2Mol-generated molecules and back to predicted expression.

We evaluated this phenotype-to-molecule-to-phenotype loop using the chemCPA-processed sci-Plex3 [33] perturbation data, focusing on 9 out-of-distribution (OOD) drugs across A549, K562, and MCF7 cells. For each cell line-drug pair, Tx2Mol-generated molecules were encoded and fed into chemCPA to predict the induced transcriptional response. The predicted response was then compared with the observed drug-induced response, defined as the difference in mean expression between perturbed and control conditions. Predictions from the matched perturbing compounds served as an upper reference (Pretrained), while those without matched drug information served as a lower reference (Baseline). Across the top 50 DEGs, Tx2Mol achieved substantially higher *R*^2^ than the Baseline and closely approached the matched-drug reference in all three cell lines (Fig. 5a). These results demonstrate that Tx2Mol-generated molecules preserve key transcriptional response information, which can be recovered by an independent perturbation predictor.

**Figure 5:**
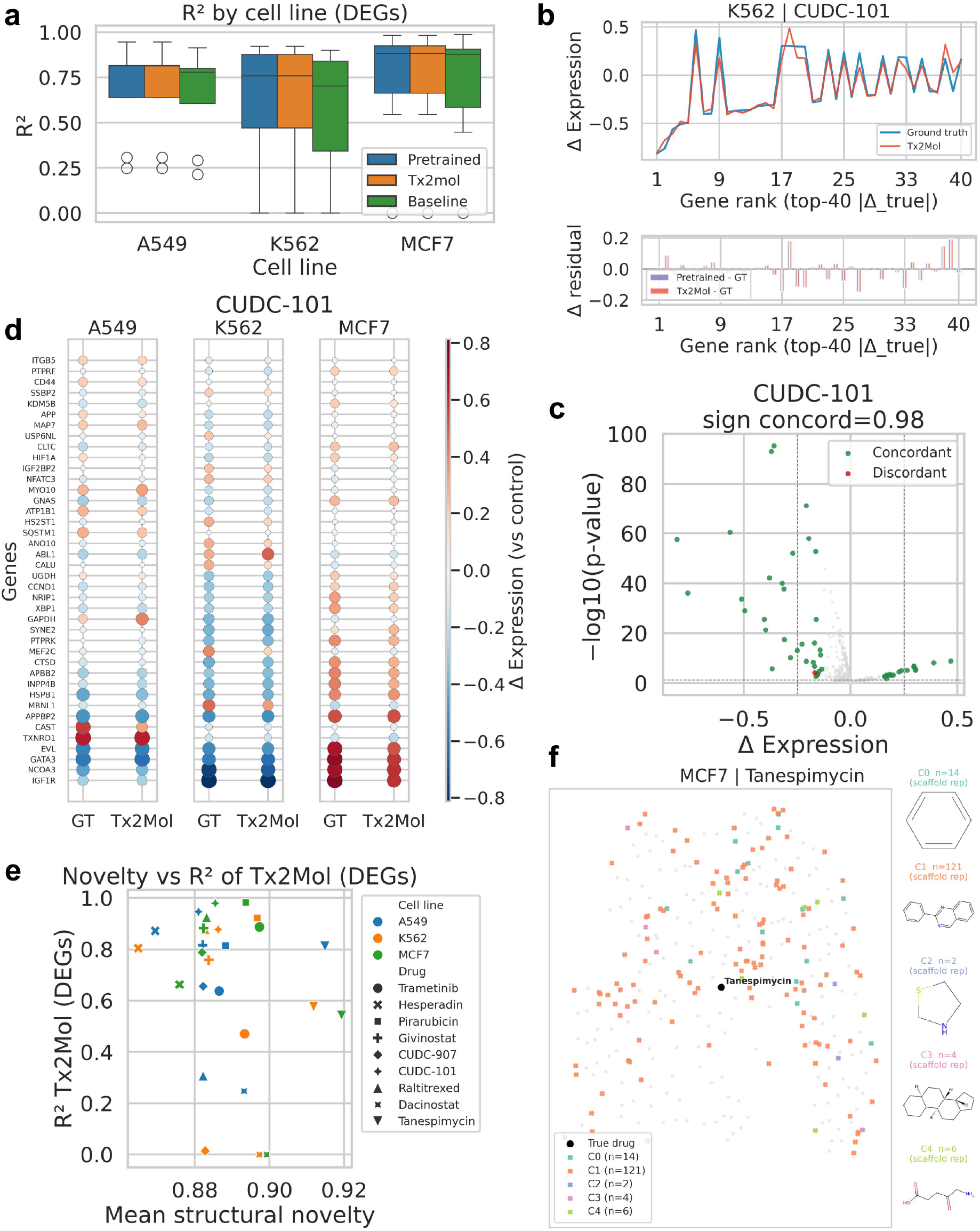
In silico validation shows that Tx2Mol-generated molecules preserve drug-induced transcriptional responses. **a**, Comparison of chemCPA-predicted and observed drug-induced responses across A549, K562, and MCF7 cells. *R*^2^ was calculated over the top 50 DEGs; Pretrained and Baseline serve as upper and lower references, respectively. **b**, Gene-level response comparison for CUDC-101 in K562 cells, showing observed and Tx2Mol-predicted Δexpression values for the top 40 response genes and corresponding residuals. **c**, Directional concordance for CUDC-101, with sign agreement computed over the top 50 genes ranked by absolute observed response. **d**, Cross-cell-line marker-gene response patterns for CUDC-101 across A549, K562, and MCF7 cells. **e**, Structural novelty versus chemCPA-predicted DEG-level *R*^2^ across cell line-drug pairs. **f**, Murcko scaffold clusters of Tx2Mol-generated molecules in the chemCPA embedding space for MCF7-Tanespimycin.

We next examined this preservation at the gene level, using CUDC-101 in K562 cells as a representative example. Tx2Mol-generated molecules reproduced the major up- and down-regulated genes among the top 40 genes ranked by the absolute magnitude of the observed response (Fig. 5b). The residuals relative to the observed response were comparable to those from the matched-drug reference. Consistently, the predicted and observed responses showed strong directional concordance, with a sign agreement of 0.98 among the top 50 response genes (Fig. 5c). Similar trends were observed across the remaining 26 cell line-drug combinations (Supplementary Figs. S8-S13). These results indicate that Tx2Mol-generated molecules preserve gene-level response patterns and the direction of drug-induced transcriptional regulation, beyond merely improving aggregate performance.

Drug responses often vary across cellular contexts. We therefore examined whether Tx2Mol preserves such context-dependent transcriptional features. Across A549, K562, and MCF7 cell lines, Tx2Mol recapitulated the top 40 marker-gene response patterns of CUDC-101, including cell-line-specific differences in up- and down-regulated genes (Fig. 5d). The predicted responses were highly consistent with the observed drug-induced response across the major marker genes. Similar cell-line-specific response patterns were observed for the remaining eight OOD drugs (Supplementary Figs. S14-S15). These results show that the predicted responses preserve cell-line-specific transcriptional signatures rather than converging to a generic drug-response profile.

We then asked whether response preservation simply reflected structural similarity to the original drugs. Across cell line-drug pairs, most Tx2Mol-generated molecules retained high DEG-level *R*^2^ despite substantial structural novelty, measured as mean 1−Tanimoto similarity to the matched real perturbing compound (Fig. 5e; Supplementary Fig. S16). This indicates that preserved transcriptional responses do not simply arise from structural imitation of existing drugs. In addition, Tx2Mol maintained high molecular validity and internal diversity across the three cell lines, with a mean validity of 0.985 and a mean diversity of 0.89 (Supplementary Fig. S17).

We further examined the scaffold organization of generated molecules using the MCF7-Tanespimycin case. Tx2Mol-generated molecules formed multiple Murcko scaffold clusters that covered the chemCPA embedding region occupied by the original drug, while also extending to nearby regions (Fig. 5f). Representative Murcko scaffolds are shown alongside the embedding, and consistent scaffold-level patterns were observed across the remaining 26 cell line-drug combinations (Supplementary Figs. S18-S24). These results provide structural support for the closed-loop analysis, showing that scaffold-diverse molecules can remain aligned with the transcriptional perturbation space of the original drug.

Together, these analyses show that Tx2Mol-generated molecules preserve drug-like transcriptional responses in silico. They recover gene-level regulatory direction, cell-line-specific response patterns, and high transcriptional fidelity despite structural novelty, supporting transcriptomic phenotypes as actionable signals for molecular design.

### 2.4 Patient-derived disease signatures guide molecular generation toward therapeutic chemical space

To evaluate whether patient-derived disease signatures can guide molecular generation, we collected 12 disease transcriptomic profiles from the CREEDS database [35]. For each disease, patient-derived gene expression changes were aggregated into a disease-level signature across 978 landmark genes. The signature was then normalized and sign-inverted to approximate a transcriptomic shift from disease toward health (Fig. 6a). Using these inverse signatures as inputs to Tx2Mol, we generated 1,000 molecules per disease and compared them with corresponding approved drugs from DrugBank [36].

**Figure 6:**
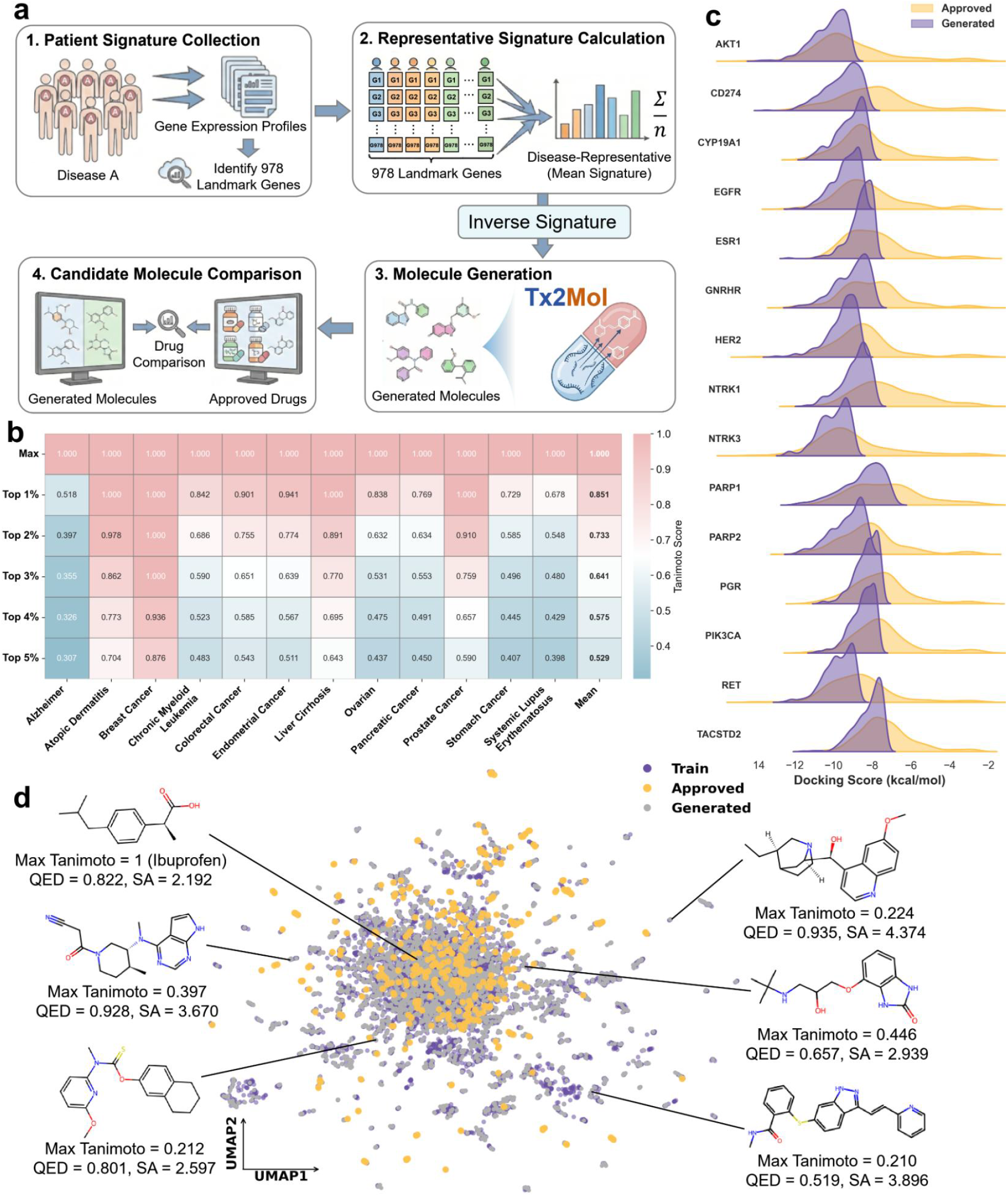
Patient-derived disease phenotypes guide molecular generation toward therapeutic chemical space. **a**, Patient-to-molecule workflow. Patient-derived expression profiles are aggregated into 978-gene disease-level signatures, sign-inverted to approximate disease-to-health shifts, and used to guide Tx2Mol generation; generated molecules are compared with approved drugs. **b**, Similarity between Tx2Mol-generated molecules and approved drugs across 12 diseases. Maximum and top 1-5% Tanimoto similarities are reported, with maximum similarity reaching 1.00 for each disease. **c**, Breast-cancer structure-based evaluation across 15 disease-relevant targets. Approved drugs are compared with the top 50% Tx2Mol-generated molecules ranked by Vina score; many candidates score below −8 kcal/mol. **d**, Chemical-space embedding of training molecules, approved drugs, and Tx2Mol-generated molecules, showing overlap with therapeutic chemical space and extension into additional chemical regions.

As shown in Fig. 6b, the maximum Tanimoto similarity reached 1.00 for each of the 12 diseases, indicating that Tx2Mol generated at least one molecule identical to a corresponding approved drug in each disease setting. Similarity enrichment remained evident among the top 1-5% of generated molecules. As progressively larger fractions of the generated set were included, similarity decreased, indicating that Tx2Mol does not collapse to near-duplicate approved-drug analogues. Instead, disease signatures guide a high-similarity subset toward approved-drug chemical space while allowing a broader exploratory tail.

We next asked whether disease-signature-guided generations also show plausible target engagement while retaining broader chemical exploration. For breast cancer, we selected 15 disease-relevant targets with high-quality structures for docking analysis (Supplementary Table S4). Approved drugs and Tx2Mol-generated molecules were docked against these targets using AutoDock Vina. The top 50% of generated molecules ranked by Vina score were selected as candidate molecules and compared with the corresponding approved drugs. These candidates showed consistently lower Vina scores across all targets, with many scoring below −8 kcal/mol (Fig. 6c), supporting strong predicted binding in silico.

Finally, a chemical embedding space map shows that Tx2Mol generations overlap regions occupied by training molecules and approved drugs while also extending into additional regions (Fig. 6d). Together, these results indicate that patient-derived transcriptomic signals can guide Tx2Mol toward approved-drug chemical space while continuing to support novel chemical exploration.

## 3 Discussion

This study addresses a central challenge in phenotype-guided molecular design: how to translate high-dimensional, noisy, and context-dependent gene-expression phenotypes into molecular structures while preserving biological guidance throughout generation. Tx2Mol was developed to maintain gene-expression guidance during molecular generation and to strengthen the link between biological responses and generated molecules. By combining persistent phenotypic guidance with transcriptome-molecule alignment, Tx2Mol shifts gene-expression signatures from passive biological readouts or one-time input conditions into actionable molecular design signals.

Across bulk gene perturbation, single-cell perturbation, and patient-derived disease settings, Tx2Mol shows that gene-expression phenotypes can guide molecular generation beyond chemical plausibility alone. In bulk perturbation benchmarks, generated molecules showed improved hit-like recovery across cancer-relevant targets while maintaining molecular quality and structural novelty. Structure-based analyses further supported plausible target engagement beyond 2D chemical similarity, as assessed by docking and ligand efficiency. In single-cell perturbation settings, Tx2Mol generalized to noisier cellular phenotypes and preserved drug-induced transcriptional responses in closed-loop in silico validation. Patient-derived disease signatures further guided generation toward approved-drug chemical space while supporting broader therapeutic chemical exploration. Together, these results connect gene-expression guidance to chemical plausibility, target relevance, and phenotype preservation, supporting phenotype-guided molecular generation as a route for early-stage hit discovery, candidate prioritization, and chemical exploration when actionable targets are unclear but disease-associated transcriptional states are measurable [37].

These findings also suggest several directions for future development. Tx2Mol provides a framework for translating gene-expression phenotypes into molecular candidates, but experimental synthesis, target-binding assays, and perturbation profiling will be needed to validate their activity. One promising direction is to extend this framework to clinically heterogeneous settings, such as drug-resistant tumors or patient-specific cancer states, where target-first discovery remains challenging but transcriptomic alterations can be measured. More broadly, drug response is shaped not only by transcriptional state but also by chromatin accessibility, proteomic context, cellular heterogeneity, and temporal dynamics of dose and exposure [38, 39]. Extending phenotype-guided generation to these richer, multimodal, and dynamic settings will require larger and better-annotated phenotype-molecule datasets, particularly those with longitudinal and multi-omics measurements [40, 41]. Future work could therefore expand Tx2Mol toward multi-omics integration, time-resolved perturbation modeling, and broader design constraints relevant to therapeutic development [42]. Coupling transcriptome-guided generation with expanded perturbation datasets, disease-specific single-cell atlases, structure-or dynamics-based screening, and iterative experimental feedback could support closed-loop drug discovery pipelines that design, test, and optimize molecular candidates directly from disease-relevant phenotypes [43, 44].

## 4 Materials and Methods

### 4.1 Overview of Tx2Mol

Tx2Mol is a transcriptome-guided post-training framework for de novo molecular generation from high-dimensional gene-expression signatures. Let *X* denote the transcriptomic perturbation space and *Y* the chemical space. Given a gene expression signature ***x*** ∈ ℝ^G^ (representing differential expression of *G* = 978 landmark genes) and an optional cell line context ***c*** as auxiliary biological information, Tx2Mol generates a SMILES token sequence ***y*** = (*y*_1_, …, *y*_*T*_) by maximizing the conditional likelihood:

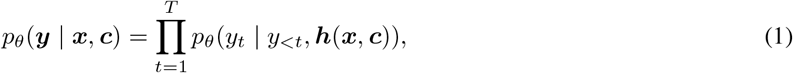

where ***h***(***x, c***) represents the projected biological guidance derived from the gene expression signature and contextual information. To couple biological guidance with molecular generation, Tx2Mol uses two complementary components: (1) **local guidance** through soft-prefix injection, which converts the biological guidance signal into learned prefix tokens that guide autoregressive decoding at each step; and (2) **global alignment** through a representation-level contrastive objective (bidirectional InfoNCE), which organizes transcriptomic and molecular representations in a shared embedding space. During post-training, the molecular backbone and conditioning projector are optimized jointly, allowing the model to learn the transcriptome-to-molecule mapping in an end-to-end manner.

### 4.2 The Tx2Mol model architecture

#### Molecular language model backbone

Tx2Mol builds on NovoMolGen-300M [45], a pretrained decoder-only Transformer for molecule de novo generation. Let *d* denote the backbone hidden size and *θ* the backbone parameters. Given an input embedding sequence, the backbone applies standard causal self-attention and feed-forward transformations to produce contextual hidden states and next-token logits for autoregressive decoding.

#### Transcriptomic latent encoder

Inspired by GxVAEs, we first pretrain a variational autoencoder (GeneVAE) on gene expression signatures to learn a low-dimensional latent representation of transcriptomic changes. Each signature is represented as a vector ***x*** ∈ ℝ^*G*^, and the encoder parameterized by *ϕ* defines an approximate posterior over latent variables 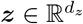:

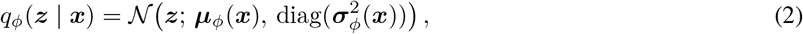

Where 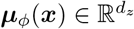 and 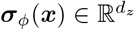 denote the encoder-predicted mean and standard deviation, respectively. We use latent dimension *d*_*z*_ = 64 and a standard normal prior *p*(***z***) = *N*(**0, *I***). GeneVAE is pretrained by maximizing the evidence lower bound [46]:

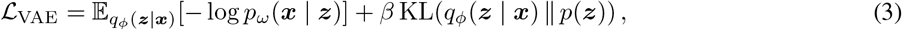

where *p*_*ω*_(***x*** | ***z***) is parameterized by the decoder network. After pretraining, GeneVAE is frozen during transcriptome-guided post-training of the molecular backbone. For downstream guidance, we use the posterior mean ***µ***_*ϕ*_(***x***) as the transcriptomic embedding ***z***. Reconstruction fidelity analyses for GeneVAE are provided in the Supplementary Figure S1.

#### Local guidance through soft-prefix

Given the transcriptomic embedding ***z*** from the frozen encoder and an optional one-hot cell-line label ***c***, we construct transcriptome-guided prefix embeddings [47] that are prepended to the molecular token stream during post-training and inference. Let ***c*** ∈ {0, 1} ^*C*^ denote the cell-line label. A learnable projection *f*_*ψ*_ maps the concatenated guidance input to *P* prefix embeddings:

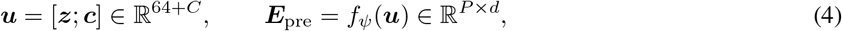

where *P* is the number of prefix tokens (default *P* = 1). Let ***E***_mol_ ∈ ℝ^*T* ×*d*^ denote the token embeddings of the SMILES sequence. The guidance-augmented input sequence is then

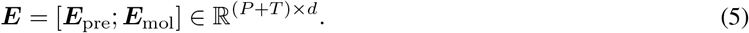

Because the prefix tokens are prepended under causal self-attention, all molecular tokens can attend to the biological guidance signal throughout decoding, enabling token-level transcriptomic guidance.

#### Global alignment through representation-level contrastive learning

In addition to token-level guidance, Tx2Mol uses a bidirectional InfoNCE objective [48] to align transcriptomic guidance signals with molecular representations at the sequence level. For each sample *i* in a mini-batch, let ***g***_*i*_ ∈ ℝ^*d*^ denote a biological-guidance representation derived from the transcriptomic signal, and let ***m***_*i*_ ∈ ℝ^*d*^ denote a molecular representation derived from the conditioned backbone. After *ℓ*_2_ normalization,

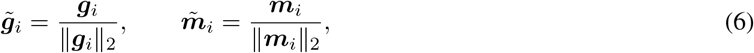

we compute, within each cell-line group *B*_*k*_, the similarity matrix

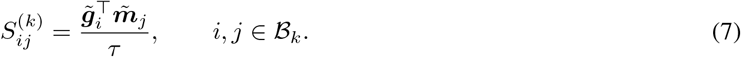

Grouping by cell line reduces trivial confounding from broad cell-context differences and encourages alignment under a comparable biological background. The group-wise contrastive loss is defined as

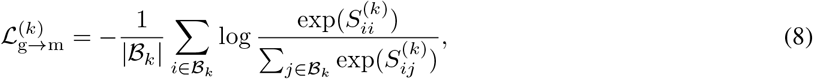

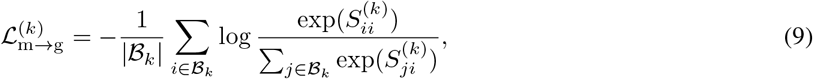

and the symmetric group-level objective is

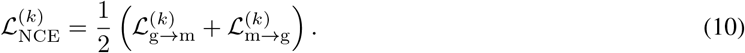

We aggregate over valid groups using a size-weighted average:

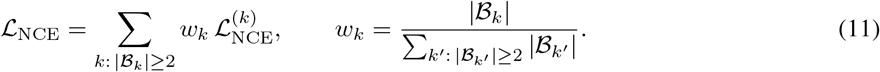

#### Joint post-training objective

Tx2Mol is post-trained by jointly optimizing a conditional language-modeling objective and the grouped contrastive alignment objective. During training, the autoregressive loss is optimized under teacher forcing. For each triple (***x, c, y***), the language-modeling loss is

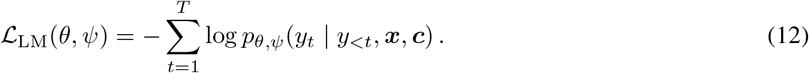

The total post-training objective is

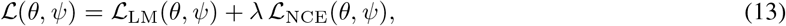

where *λ* controls the contribution of the contrastive term. During post-training, all backbone parameters *θ* and conditioner parameters *ψ* are updated, while the transcriptomic encoder parameters *ϕ* remain frozen.

### 4.3 Guided generation at inference

Given a new gene-expression signature ***x***^*^ and a designated cell-line label ***c***^*^, Tx2Mol first passes ***x***^*^ through the frozen GeneVAE encoder to obtain a transcriptomic embedding ***z***^*^. The embedding ***z***^*^ is then concatenated with the corresponding cell-line context ***c***^*^ and mapped by the learned guidance projector to a sequence of prefix embeddings 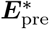. These prefix embeddings are prepended to the molecular language model input and retained throughout autoregressive decoding. Guided by 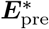, the backbone generates SMILES tokens sequentially. For settings without an intrinsic cell-line context, including patient-derived disease signatures, we used the most frequent post-training cell-line context (MCF7) as the default auxiliary context.

To generate multiple candidates for each guidance input, we used stochastic decoding with temperature scaling (*T* = 1.0), top-*p* nucleus sampling (*p* = 0.95), and top-*k* truncation (*k* = 100). Unless otherwise specified, 100 molecules were generated per guidance input, with a maximum SMILES length of 100.

### 4.4 Data collection and preprocessing

Tx2Mol was evaluated across three settings: bulk gene perturbation signatures, single-cell perturbation signatures, and patient-derived disease signatures. To enable a shared guidance scheme across data sources, all transcriptomic inputs were represented as a fixed-length gene-expression change vector ***x*** in the 978-dimensional landmark-gene space, optionally accompanied by a cell-line label ***c*** as contextual information.

#### Post-training dataset

Post-training was performed using bulk transcriptome-molecule pairs derived from preprocessed LINCS data, comprising more than 3 × 10^4^ paired samples 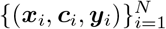, where ***y***_*i*_ denotes the SMILES string associated with transcriptomic signature ***x***_*i*_ in cell line ***c***_*i*_. We retained compound-treated samples measured at 10 *µ*M for 24 h and grouped them by cell line-compound pair. For each group, we calculated the average pairwise Pearson correlation (APC) among samples within the same cell line-compound pair; groups with APC < 0.7 were considered unreliable and excluded. For the remaining groups, gene-expression signatures were aggregated by the median to obtain a single profile per cell line-compound pair. Finally, we retained the 14 cell lines with the largest number of samples, and split the data within each cell line into training and validation sets at a 9:1 ratio.

#### Bulk gene perturbation signatures for evaluation

To assess hit-like generation under genetically perturbed transcriptional states, we collected target-gene perturbation profiles from the LINCS database [49]. We focused on ten proteins with established relevance as therapeutic targets in cancer: AKT1, AKT2, AURKB, CTSK, EGFR, HDAC1, MTOR, PIK3CA, SMAD3, and TP53. For the first eight proteins, transcriptomic signatures were derived from gene knockdown experiments in the MCF7 cell line; for the PIK3CA and TP53 proteins, signatures were derived from gene overexpression experiments. When multiple profiles were available for the same target under different experimental conditions, they were averaged to obtain a single target-level perturbation signature.

#### Single-cell perturbation dataset for evaluation

To evaluate performance on single-cell perturbation data, we used the sci-Plex 3 dataset and retained three cell lines: A549, MCF7, and K562. Only human genes were kept, and gene features were restricted to the intersection with the 978 LINCS landmark genes. At the cell level, we retained single-drug perturbation samples measured at 10 *µ*M for 24 h, and excluded cells associated with combinatorial perturbations or missing compound annotations. Gene expression values were normalized using log(1 + *X*). PCA and Harmony [50] were used only for preprocessing and downstream visualization, not as model inputs. To define transcriptomic guidance signals for generation, samples were grouped by cell line-compound label, and cells within each group were aggregated by the median to obtain a single transcriptomic profile in the 978-gene landmark space. These aggregated profiles were used as transcriptomic guidance signals for generation and downstream evaluation.

#### Patient-derived transcriptomic profiles for disease-signature-guided generation

Patient-derived transcriptomic profiles were used only for generation and downstream evaluation, and were not included in model training or parameter updates, thereby serving as an out-of-distribution evaluation setting. For this analysis, we obtained disease-associated expression profiles from the Crowd Extracted Expression of Differential Signatures (CREEDS) database, which contains transcriptomic data for 79 diseases. We selected 12 diseases: Alzheimer’s disease, atopic dermatitis, breast cancer, chronic myeloid leukemia, colorectal cancer, endometrial cancer, liver cirrhosis, ovarian cancer, pancreatic cancer, prostate cancer, stomach cancer, and systemic lupus erythematosus. For each disease, we restricted the profiles to the overlap with the 978 LINCS landmark genes and averaged expression changes across patients to obtain a disease-level signature. The resulting signatures were sign-inverted to approximate gene-expression changes associated with a transcriptional shift from disease toward health. Approved drugs used for downstream comparison were obtained from DrugBank.

### 4.5 Evaluation protocol

We evaluated Tx2Mol across three validation dimensions: chemical plausibility, structural compatibility, and phenotypic preservation.

#### Chemical plausibility

Generated molecules were evaluated for validity, uniqueness, and novelty after SMILES canonicalization with RDKit. We quantified validity as the fraction of chemically valid molecules among all generations, uniqueness as the fraction of non-duplicate molecules after canonicalization, and novelty as the fraction of generated molecules absent from the post-training set. Molecular quality was further characterized by QED, SA, LogP, and MW, which summarize overall drug-likeness, synthetic accessibility, hydrophobicity, and molecular size, respectively. Chemical-space relationships among generated molecules, training molecules, and approved drugs were visualized using UMAP projections of Morgan fingerprints. Structural novelty beyond exact overlap was quantified by a structural novelty index (SNI),

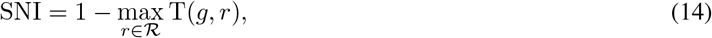

where *g* denotes a generated molecule, R the reference set of approved drugs from DrugBank (U.S., Canada, and E.U.), and T(·, ·) the Tanimoto similarity. Larger SNI values indicate greater structural novelty relative to approved drugs.

#### Structural compatibility

To assess binding plausibility beyond 2D similarity, generated molecules were docked into experimentally resolved target pockets using AutoDock Vina. Target structures and ligand-bound pockets were curated from experimentally resolved target-ligand structures annotated through UniProt/PDB resources, and docking analyses were performed for both bulk target benchmarks and disease-level evaluations. Lower Vina scores were interpreted as stronger predicted binding. Because raw docking scores can favor larger molecules, we additionally computed ligand efficiency (LE) [51] to normalize predicted binding by molecular size:

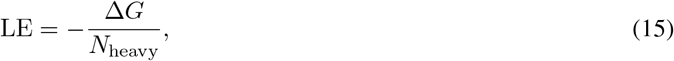

where Δ*G* is approximated by the Vina score and *N*_heavy_ is the number of heavy atoms in the molecule. LE was reported together with the Vina score to account for molecule size when comparing predicted binding strength.

#### Phenotypic preservation

Phenotypic preservation and condition specificity were assessed using setting-specific reference comparisons. For a generated molecule *g* and a reference molecule *r*, Tanimoto similarity was defined as

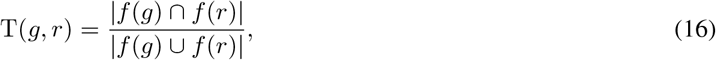

where *f* (·) denotes the corresponding Morgan fingerprint. In the bulk gene perturbation benchmark, generated molecules were compared with the corresponding known ligands, and cross-target specificity was further assessed by comparing each on-target generation set against known ligand sets from all targets. On-target and off-target similarity were additionally compared using cosine similarity to test whether generated molecules preferentially aligned with the intended perturbation condition. In the single-cell setting, phenotype consistency was further evaluated with a compound-phenotype compatibility scorer trained on true transcriptome-molecule pairs and hard negatives, with performance summarized by AUROC, AUPRC, and score distributions. Pathway-level analyses were used to examine whether molecules generated under the same guidance signal remained phenotypically coherent while preserving chemical diversity.

#### In silico drug-perturbation validation

To evaluate whether Tx2Mol-generated molecules preserve drug-induced transcriptional responses, we used chemCPA [34] as an independent in silico perturbation-response predictor. For each cell line-drug pair in the chemCPA-processed sci-Plex3 dataset, Tx2Mol-generated molecules were encoded by the chemCPA molecular encoder and used to predict the induced transcriptional response. Predicted responses were compared with the observed drug-induced response, defined as the difference in mean expression between perturbed and control cells. Predictions conditioned on the matched perturbing compound were used as an upper reference, whereas predictions without matched drug information were used as a lower reference. Response preservation was quantified using *R*^2^ over the top 50 differentially expressed genes ranked by the absolute observed response, gene-level residuals, directional sign agreement, and cell-line-specific marker-gene response patterns.

## 5 Data availability

All datasets analysed in this study were obtained from publicly available resources. Bulk compound-perturbation and gene-perturbation transcriptomic data were obtained from LINCS (https://clue.io/data/CMap2020#LINCS2020); the sci-Plex 3 dataset was downloaded from GEO under accession GSM4150378 (https://www.ncbi.nlm.nih.gov/geo/query/acc.cgi?acc=GSM4150378); patient-derived disease expression signatures were obtained from CREEDS (https://amp.pharm.mssm.edu/creeds/); approved-drug information was obtained from DrugBank (https://go.drugbank.com/); and protein structures used for docking analyses were identified via UniProt and obtained as PDB files (https://www.uniprot.org/; https://www.rcsb.org/).

## 6 Code availability

The code used in this study is available at https://github.com/AI-HPC-Research-Team/Tx2Mol.

## Acknowledgement

This work was supported by Guangdong S&T Programme (Grant No.2024B0101010003). The authors thank Peng Cheng Cloud-Brain for the valuable support during this research. We gratefully acknowledge computational support from Shanghai Smart Logic Technology and Tianqiong supercomputing platform.

